# Chiral Derivatization-enabled Discrimination and Visualization of Proteinogenic Amino Acids by Ion Mobility Mass Spectrometry

**DOI:** 10.1101/2022.07.04.498692

**Authors:** Chengyi Xie, Yanyan Chen, Xiaoxiao Wang, Yuanyuan Song, Yuting Shen, Xin Diao, Lin Zhu, Jianing Wang, Zongwei Cai

## Abstract

The importance of chiral amino acids (AAs) in living organisms has been widely recognized since the discovery of endogenous D-AAs as potential biomarkers in several metabolic disorders. Chiral analysis by ion mobility spectrometry-mass spectrometry (IMS-MS) has the advantages of high speed and sensitivity but is still in its infancy. Here, a N_α_-(2,4-dinitro-5-fluorophenyl)-L-alaninamide (FDAA) derivatization is combined with trapped ion mobility spectrometry-mass spectrometry (TIMS-MS) for chiral AA analysis. For the first time, we demonstrate the simultaneous separation of 19 pairs of chiral proteinogenic AAs in a single fixed condition TIMS-MS run. The utility of this approach presents for mouse brain extracts by direct-infusion TIMS-MS. The robust separation ability in complex biological sample was proven in MALDI TIMS mass spectrometry imaging (MSI) as well by directly depositing 19 pairs of AAs on a tissue slide following on-tissue derivatization. In addition, endogenous chiral amino acids were also detected and distinguished. The developed methods show compelling application prospects in biomarker discovery and biological research.

**Entry for the Table of Contents:** 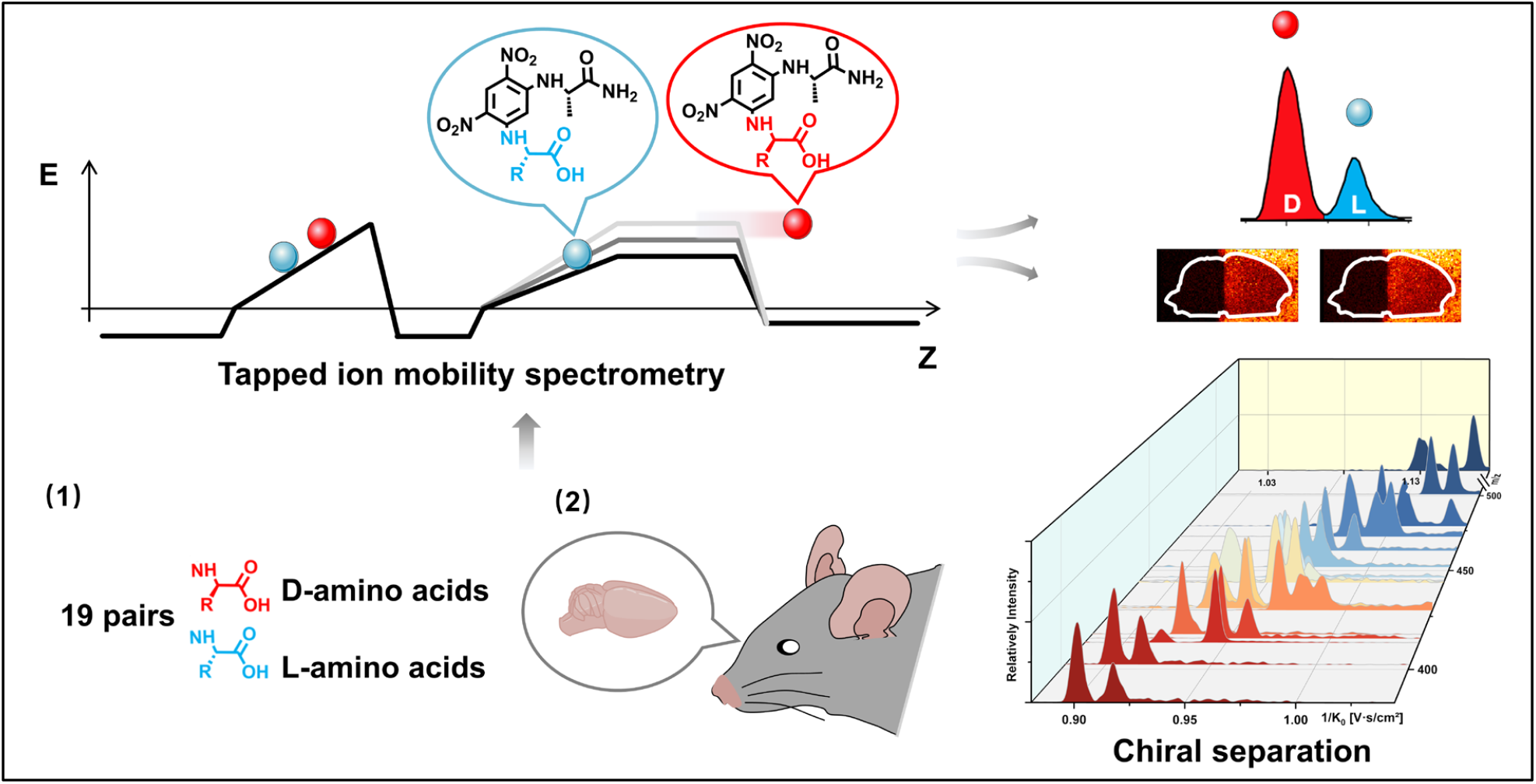

The combination of chiral derivatization and trapped ion mobility-mass spectrometry provides the first insights into the separation of 19 pairs of chiral proteinogenic D/L-amino acids in a single run and further visualization of chiral amino acids under complex biological matrix.

## Introduction

Chiral asymmetry is ubiquitous in nature. In organisms, an asymmetric feature is that amino acids (AAs) are basically in L-form, but only a few D-AAs appear. Free D-AAs and related metabolites have recently drawn increasing attention since their important roles in physiological activities and neurological disorders^1^. It was found that D-Ser, generated from the racemization of L-serine in mammal’s brains^2^, is a coagonist of the N-methyl-D-aspartate (NMDA) receptors^3^. In addition, the abnormal level of D-Ser has been associated with neurological diseases, such as Alzheimer’s disease^4^ and schizophrenia^5^. The D-Ser was found in organisms other than mammals as well. It was reported that D-Ser plays a role in sleeping regulation in *Drosophila* intestine^6^ and is related to epidermal vesicle release in chordate Ciona^7^. However, the involvement of most D-AAs in physiological activities is still not clearly understood. Furthermore, D- and L-AAs have highly similar chemical and physical properties and differ only in their optical activity, hindering their routine separation for analysis and further biofunctional studies in complex samples. Chiral research highly relies on the development of analytical methodologies with adequate selectivity and sensitivity to achieve the detection of low concentrations of D-AAs from the interference of abundant L-AAs in biological samples.

Electrophoretic and chromatographic techniques have been widely used for chiral analysis^8^. In general, chiral stationary phase, chiral selectors or chiral derivatization agents (CDA) are involved in the separation process and have been well developed in recent years^9–12^, promoting the depth and wideness of chiral research. The combination of mass spectrometry (MS) and capillary electrophoresis (CE) or liquid chromatography (LC) can further improve the performance of chiral identification because the mass spectrometry can accurately determine molecular weight and provide molecular fragmentation information. Even so, the time-consuming nature limits further applications of liquid-phase separation techniques for rapid chiral assay and in-situ/imaging analysis.

Ion mobility spectrometry (IMS) largely compensates for the drawback of CE and LC as it provides gas-phase separation on a millisecond time scale compared to the minutes to hours separation in CE or LC. In addition to isomeric separation^13–16^, the fast speed IMS can be coupled with the liquid phase separation to greatly increase the peak capacity and dynamic range by constituting a multidimensional separation system^17^. IMS-MS has been used for the investigation of the chiral effects of truncated amyloid-beta (Aβ) related to Alzheimer’s disease^18^. Recently, it has been reported that protonation-induced diastereomers can be separated by differential mobility spectrometry (DMS)^19^. In IMS, gas-phase ions are separated according to their average collisional cross section (CCS) by the combinational manipulations of the electrical field and inert-gas flow^20^. Nevertheless, enantiomers could not be separated directly by IMS since they have the same mobility in the gas phase ^21,22^. Therefore, different methodologies, (i) the introduction of volatile chiral reagent into the IMS drift tube^23^, (ii) formation of noncovalent complexes ^21,24–28^, and (iii) chemical derivatization methods^22,29,30^, have been developed for chiral IMS analysis. In general, the core principle of methods (ii) and (iii) is transforming enantiomers into diastereomers to generate CCS differences between pairs of original enantiomers in IMS. The success of the gas-phase chiral separation strategy was made possible by the development of high resolving power (R) IMS instrumentation, such as trapped ion mobility spectrometry (TIMS)^31^, cyclic ion mobility (cIM)^32^, structures for lossless ion manipulations (SLIM)^33^, U-shaped mobility analyzer (UMA)^34^, DMS^35^, and so on. However, the highly similar chemical structure limits further separation of small chiral amino acids even with high *R* IMS^29^.

With the combination of high resolving power IMS and novel chiral derivatization methods, continuous progress has been made for direct chiral IMS with the increasing number of chiral AAs to be separated using a single chiral selector^21,28,29^. In 2019, Simon et al. resolved 13 proteinogenic AAs using chiral derivatization with (+)-1-(9-fluorenyl)ethyl chloroformate (FLEC)^22^. Karst et al. developed an automated method for the separation of eight AAs by chiral derivatization with (S)-naproxen chloride (S-NAP) and following TIMS-MS analysis^29^. It should be noted that by optimizing the IMS conditions individually for each pair of amino acids, Guo et al. achieved the separation of all 19 proteinogenic AAs using a steroid-based chiral derivatization reagent by a U-shaped ion mobility-mass spectrometer^30^. However, individually optimized IMS conditions may limit the application of this method for high-throughput chiral analysis and cannot be applied to *in situ* imaging analysis. In the view above, the development of chiral derivatization methods with fixed IMS conditions to simultaneously separate all proteinogenic AAs remains an analytical challenge. The significance of achieving this goal lies in the direct detection and practical application of IMS-based chiral selectors in complex biological samples, including a large number of isomeric and isobaric interference. Furthermore, the relative abundance of chiral AA has been observed to vary across different areas in brain tissue^4^, providing a strong motivation to visualize chiral AA distributions by mass spectrometry imaging (MSI).

In this work, the combination of N_α_-(2,4-dinitro-5-fluorophenyl)-L-alaninamide (FDAA) derivatization and TIMS-MS for chiral amino acid analysis was systematically investigated. The powerful IMS-based chiral separation ability of FDAA was demonstrated for all 19 proteinogenic AAs, using the same ionic form without tedious optimization. To the best of our knowledge, FDAA derivatization achieved for the first time the successful differentiation of the mixture of 19 pairs of DL-AAs with concentration down to nM in a single TIMS-MS run and the acquisition time of about one minute. Moreover, the effectiveness of FDAA for AA identification was also investigated in mouse brain samples.

## Experimental

### Chemicals and samples

D and L forms of arginine (Arg), alanine (Ala), tryptophan (Trp), leucine (Leu), aspartic acid (Asp), glutamine (Gln), glutamic acid (Glu), phenylalanine (Phe), proline (Pro) as well as D forms of asparagine (Asn), histidine (His), methionine (Met), serine (Ser), threonine (Thr), tyrosine (Tyr), valine (Val), and L form of isoleucine (Ile) were purchased from Aladdin Co., Ltd. (Shanghai, China). L-His, D-lysine (Lys), L-Lys, L-Met, L-Thr, L-Val, L-Ser, N_α_-(2,4-dinitro-5-fluorophenyl)-L-alaninamide (FDAA), N_α_-(2,4-dinitro-5-fluorophenyl)-L-valinamide (FDVA), sodium bicarbonate, HPLC-grade tetrahydrofuran (THF), 2,5-Dihydroxyacetophenone (DHAP) were obtained from Sigma Aldrich Corp. (Darmstadt, Germany). D-Cysteine (Cys), L-Asn, and D-Ile were purchased from Shanghai Macklin Biochemical Co., Ltd. (Shanghai, China). L-Cys was obtained from Shanghai Yuanye Biological Technology Co., Ltd. (Shanghai, China). L-Tyr was purchased from Beijing Dingguo Changsheng Biotechnology Co., Ltd. (Beijing, China). HPLC-grade methanol (MeOH) was obtained from VWR Chemicals (Fontenay-sous-Bois, France). AR-grade chloroform was purchased from RCI Labscan (Bangkok, Thailand). Pesticide-grade acetone was purchased from Duksan Pure Chemical Co. Ltd. (Ansan, Kyonggido, South Korea). Deionized water was produced from a Milli-Q water purification system (Millipore Corp., Bedford, MA, USA).

### Sample preparation

Amino acid standard solutions were prepared separately by dissolving solute into deionized water at the concentration of 1 mM except for Asp and Glu solutions (10mM) which use aqueous NaOH (1 mM) as a solvent. D/L-AA mixture solutions were prepared by mixing D- and L-AA at a 1:1 volume ratio. Similarly, the mixture solution of 38 AAs was prepared by mixing all AA standards at the same volume. For the determination of enantiomeric ratio in AA mixtures, D-AA and L-AA with different volume ratios (0.5/99.5, 1/99, 10/90, 20/80, 30/70, 40/60, 50/50) were prepared. The preparation of mouse brain extracts and tissue slides are provided in the Supporting Information. For chiral derivatization, equal volumes (20 *µ*L) of the prepared analyte solution (Pure AA solution, mixed AA solution, or brain extract), aqueous sodium bicarbonate (100 mM), and derivatization reagent (FDAA or FDVA, 5 mM) dissolved in MeOH/THF (1:1, v/v) were fully mixed and then incubated at 40 °C for 60 minutes. For the analysis of AA standards, all derivatized mixtures were diluted 100 times using methanol. Therefore, the final concentrations for each AA were 3.3 *µ*M, 1.65 *µ*M, and 87.7 nM except Asp and Glu (33 *µ*M, 165 *µ*M, and 877 nM) in the pure standards, racemic mixtures, and 38 AA mixtures, respectively. For the analysis of mouse brain extracts, the derivatized mixtures were diluted 10 folds in methanol before the direct-infusion TIMS-MS analysis.

### Direct-infusion TIMS-MS

All ESI TIMS-MS and MALDI TIMS-MS were performed on timsTOF fleX MALDI-2 (Bruker Daltonics, Bremen, Germany) equipped with dual ESI-MALDI sources. The ESI source was used for direct-infusion study with a flow rate of 3 *µ*L/min and the parameters of the ion source were set as the following: end plate offset, 500 V; capillary voltage, 4000 V; nebulizer, 0.3 bar; dry gas, a flow rate of 3.5 L/min at 200 °C. For the TIMS analyzer, the accumulation time was set to 100 ms. 1/*K*_0_ range (0.7−1.2 V·s/cm^2^) and ramp time (949 ms), which mainly determine the resolving power of TIMS, were kept constant throughout all direct-infusion experiments. The mass range was set to 150-1300 m/z to include all analyte ions. In addition, parameters in Tune are listed in Table S1, which influence the detection range and sensitivity of TIMS-MS as well.

## MALDI TIMS-MS

Data were acquired at 100 *µ*m spatial resolution in M5 mode with 100% laser power at 1000 Hz and 200 shots per pixel. The ramp time and 1/*K*_0_ range were adjusted to 839 ms and 0.7−1.3 V·s/cm^2^. The accumulation time was 200 ms, which was determined by laser shots and frequency in the MALDI mode. Other parameters were listed in Table S2.

### Calibration

Mass and mobility calibrations were performed before the start of each experiment using the ESI-L Low Concentration Tuning Mix (Agilent Technologies, CA, USA). Since the elution voltage *V*_*e*_ is tightly correlated to the reduced mobility of the ions^36^, the calibration curve is derived from *K*_0_ = *a* + *b*/*V*_*e*_, where *a* and *b* are calibration constant and were determined from known calibrant ions in the tuning mix. Before the TIMS calibration, the gas flow was fine-tuned to ensure that *V*_*e*_ value of the calibrant ion at m/z 622.03 is within 132±1 V. The calibration process was accomplished by timsControl (Bruker Daltonics, Bremen, Germany). Then the reduced mobility of unknown analytes can be determined as long as the TIMS operating conditions remain to be constant.

### Data analysis

The extraction of m/z, ion mobilograms, 1/*K*_0_, and resolving power (*R*) and finding of mobilogram compounds were accomplished by Compass DataAnalysis 5.3 (Bruker Daltonics, Bremen, Germany). MALDI-TIMS MS data was normalized on the root mean square value (RMS) and visualized using SCILS Lab MVS (2022b Premium 3D, Bruker Daltonics, Bremen, Germany). IMS resolution (*R*_*pp*_) and *R* were calculated as *R*_*pp*_ = 1.18(A2 − A1)/(ΔA1 + ΔA2) and *R*= A/ΔA, respectively, where A is the 1/*K*_0_ value and ΔA is full peak width at half-maximum.

## Results and discussion

### Single condition differentiation of nineteen chiral amino acid pairs

The differentiation of enantiomers is the prerequisite for further exploring the roles and functions of chiral amino acids in living organisms. Although continuous progress has been made for IMS-based chiral separation^21–26,28–30^, the challenge of the realization of a single IMS separation analysis for a large number of AAs in complex samples has not been overcome due to the requirement of not only high sensitivity for trace levels of chiral analytes but also the ability to distinguish enantiomers under the interference of other isobars and isomers. Given the proven potential of chiral derivatization agent N_α_-(2,4-dinitro-5-fluorophenyl)-L-alaninamide (FDAA) in chiral LC analysis^37–40^, we used FDAA derivatization for IMS-based chiral separation in both standards and real samples. The derivatization reaction scheme is shown in Fig.1A. Nucleophilic aromatic substitution of FDAA by the amino group of amino acids leads to the transformation of enantiomers into diastereomers^37^. To clarify the chiral-IMS analysis process, the standard solution of D-Ser was selected as a representative. Fig.1B shows the mass spectrum of the derivatized D-Ser solution after the direct-infusion analysis. Due to the use of sodium bicarbonate buffer in the reaction mixtures, [FDAA-D-Ser + 2Na – H]^+^ is observed at *m/z* 402.063 in the mass spectrum. According to previous studies, the formation of metal ion adducts facilitates the differentiation of chiral amino acids^21,22,24,25,29,30,41^, especially the [M + 2Na - H]^+^ ionic form of the AAs’ derivatives^22,30^. Since diastereomers have exactly the same molecular weight, their separation is achieved exclusively by IMS. The ion mobilogram of *m/z* 402.063 ± 0.01 was extracted by plotting 1/*K*_0_ (inverse reduced mobility; To simplify, the term “mobility” will be used) against intensity to compare the mobility distribution of the formed diastereomers (Fig.1B lower panel).

**Fig.1.**
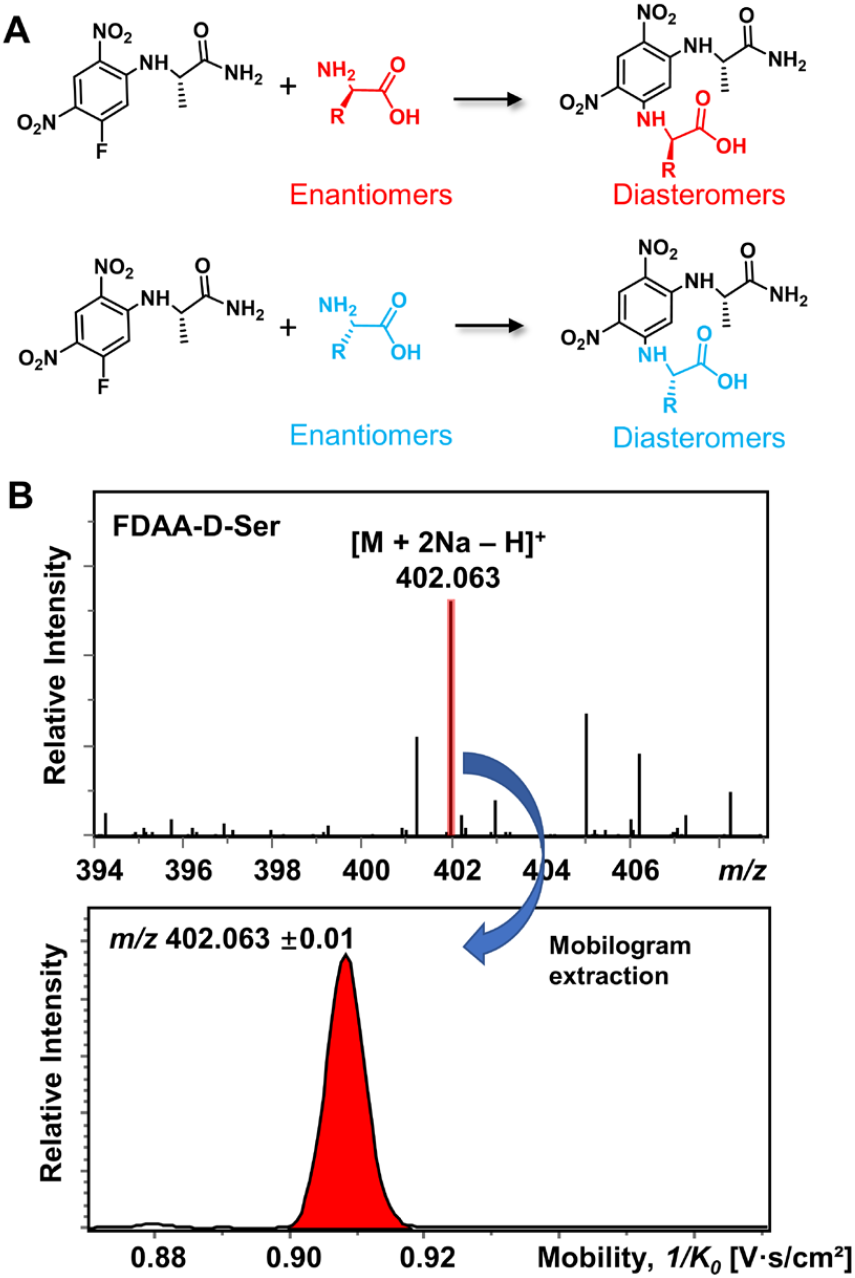
(A) Chiral derivatization reaction of AAs using FDAA. (B) Mass spectrum of FDAA derivatized Ser and the corresponding extracted mobilogram at *m/z* 402.063 ± 0.01.

All 19 proteinogenic amino acids were investigated to evaluate the effectiveness of the FDAA derivatization and chiral separation by TIMS-MS. The extracted ion mobilograms (EIMs) of 19 pairs of FDAA-AA mixtures are presented in Fig.2A-2S. All the measurements of TIMS-MS experiments were carried out under the same instrumental settings (same TIMS and Tuning parameters), enabling comparison between different pairs of diastereomers and reducing tedious optimization processes for each set of experiments. To keep the same instrumental settings, the mobility range should be set large enough to include all analytical ions while the separation performance of TIMS will be compromised. In order to quantitatively compare separation degree, the distribution of peak-to-peak resolution *R*_*pp*_ is shown in Fig.2T. In addition to D/L mixtures, the EIMs of their pure standards (i.e., D-AA and L-AA measured separately) are shown in Fig.S1. The result confirmed that our method achieves effective separation on D/L AAs and eliminates the possibility of multiple mobility peaks caused by other factors (i.e. different ionization sites ^42–45^ or structural dynamics during transit through TIMS^46^). In general, all 19 pairs of chiral AAs could be distinguished by only selecting the ionic form of [M + 2Na – H]^+^ without carefully optimizing the combination of reference compounds and cation ions. Among them, baseline or nearly baseline separation could be achieved for most chiral AAs, which is consistent with the observation that most *R*_*pp*_ values shown in Fig.2T are over one. All these results strongly demonstrate that FDAA is a universal CDA for the analysis of chiral AAs by TIMS. For D/L mixtures, the mobility distribution of their separated peak in Fig.2A-2S can be aligned to their pure enantiomeric standards separately measured in Fig.S1, indicating FDAA-AA diastereomers are indeed separated by TIMS owing to the considerable structural difference between FDAA-AA diastereomers in the gas-phase. On the other hand, the use of a fixed separation condition enables the maintenance of stable separation results (e. g., 1/*K*_0_ and *R*_*pp*_) for various samples (e. g., individuals and mixtures), facilitating the accurate and effecient identification of targeted analytes in real samples without optimization case by case.

**Fig.2.**
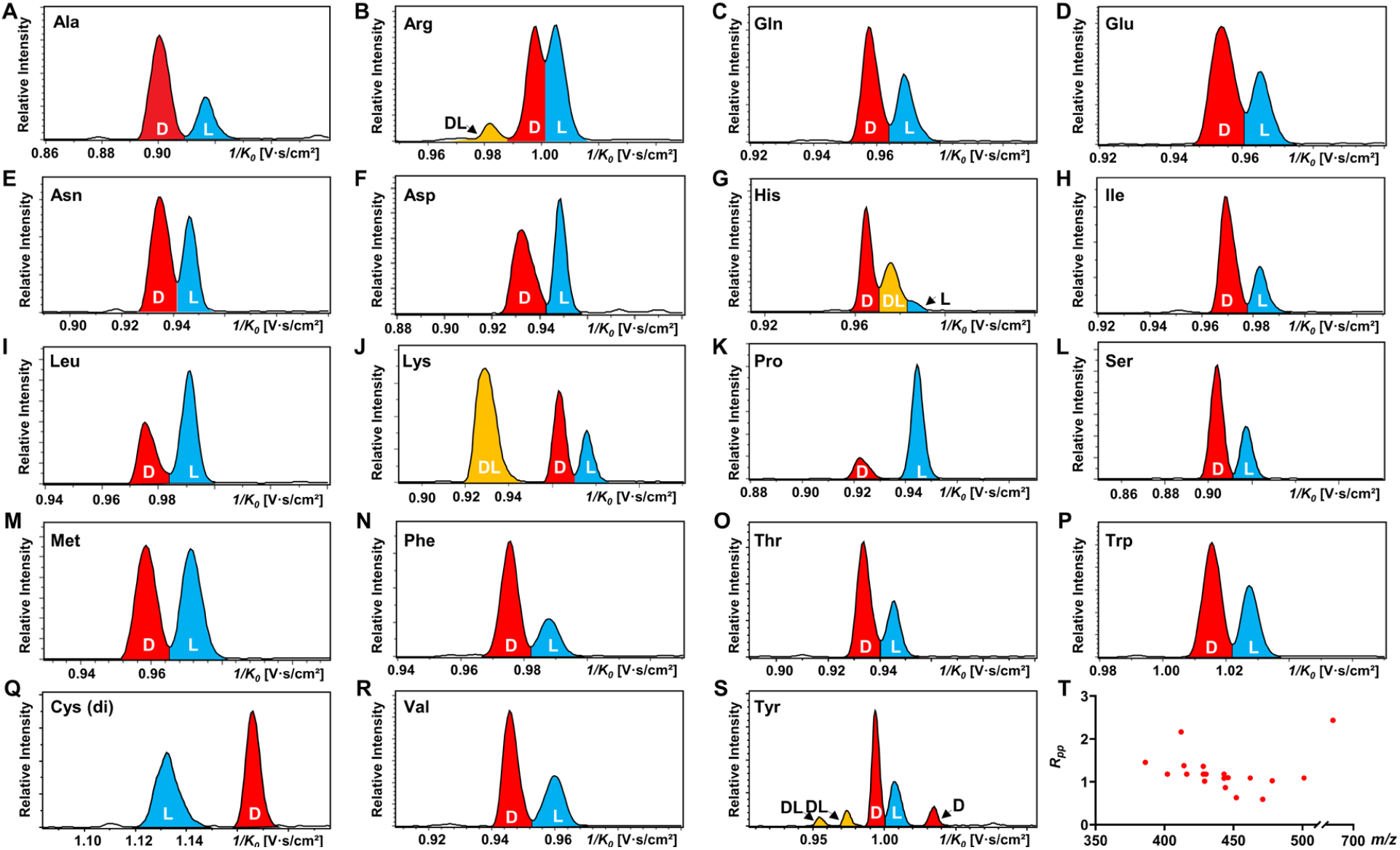
TIMS differentiation of 19 chiral amino acid pairs under constant separation conditions without individual optimization for each amino acid. (A) to (S) EIMs of AAs (D: L = 1: 1) after FDAA derivatization, where peaks in red, blue, and yellow traces represent D-, L-, and DL-AAs, respectively. Cys (di) indicates bis-derivative of Cys. (T) Peak-to-peak resolution (*R*_*pp*_) distributions of resolved chiral AAs in Fig.2A - 2S. In the case of multiple peaks, only the smallest *R*_*pp*_ value is considered.

Another chiral derivatization agent analogous to Marfey’s reagent, N_α_-(2,4-dinitro-5-fluorophenyl)-L-valinamide (FDVA), was induced in this study to compare the differentiation capabilities of FDAA and FDVA for chiral amino acids. TIMS analysis for each pair of AA mixtures after FDVA derivatization was performed under the same instrumental settings as the FDAA method, and the corresponding EIMs are presented in Fig.S2. The FDAA derivatization method showed a generally better separation ability than FDVA for chiral AAs. All AAs are resolved by FDAA derivatization, while three AAs (Ala, Arg, and Thr) could not be resolved by FDVA derivatization. The above results indicated that FDAA is a better choice for the chiral IMS analysis of amino acids.

In addition, multiple features of Arg, His, Lys, and Tyr are observed in the EIMs of AA mixtures in Fig.2 and pure optical standards in Fig.S1 after FDAA derivatization. In the case of basic amino acids, Arg and Lys contain two primary amines, leading to different positional substitutions by FDAA. Moreover, FDAA can also react with thiols and aromatic alcohols^38^, resulting in multiple peaks in the mobility dimension for Tyr and bis-derivatives for Cys. The arising of multiple peaks is consistent with the chromatographic separation of FDAA derivatized AAs^38^. Even though the double substitutions increase the complexity of chiral analysis, unique features for each chiral AA can be resolved from its optical isomer by the TIMS analyzer.

The observed elution order and resolving power *R* of the resolved main peaks in Fig.2 are listed in Table S3. Regarding the elution order, derivatized D-AAs generally elute first from the TIMS tunnel than their L-form counterparts, indicating a more extended gas-phase structure (higher reverse reduced mobility) of the former. The only observed exception is the bis-derivatives of D- and L-Cys, which shows the opposite elution order (Fig.2Q). The resolving power listed in Table S3 basically ranges from 100 to 200. Since the resolving power of TIMS also related to the *m/z* of analytes^47^, in addition to the parameter settings of the TIMS analyzer, a slight shift of *R*values is observed for different AA derivatives. Considering the resolving power of about 200 already achieved by other IMS instrumentations (e.g. UMA^34^, cIM^32^, DMS^35^, and SLIM^33^), chiral analysis of AAs by FDAA derivatization can transfer to other IMS platforms without a further improvement in sample pretreatment methods.

### IMS-MS differentiation of all proteinogenic chiral amino acids mixture in a single TIMS-MS run

To our knowledge, chiral separation of all proteinogenic amino acids by a single fixed IMS-MS condition has not been achieved. Based on the results described above in this study, we further attempted to realize the possibility of distinguishing all 19 pair chiral AA mixture in a single TIMS-MS run. The stacking EIMs of the derivatized AAs are shown in Fig.3A arranged from front to back according to their *m/z* values observed in the mass spectrum (Fig.S3). Separately listed mobilograms are shown in Fig.S4 to provide a clearer visualization of Fig.3A.

**Fig.3.**
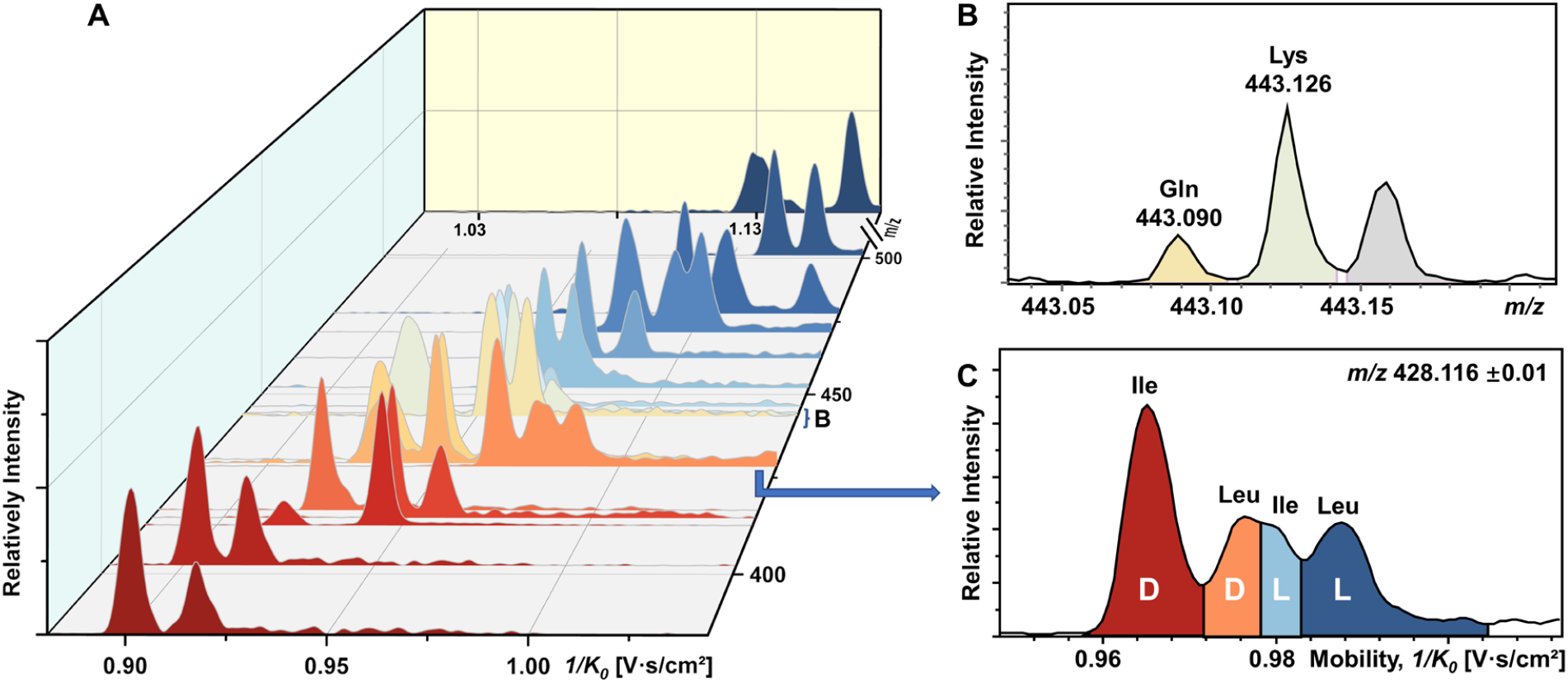
Direct infusion analysis of 38 mixed amino acids by FDAA derivatization in a single TIMS-MS run. (A) Stacking EIMs of 38 chiral AAs obtained from the mixture. (B) Mass spectrum of fully resolved isobars (Gln and Lys). (C) Mobilogram extracted from *m/z* 428.116 ± 0.01 containing four resolved isomers (D-Ile, D-Leu, L-Ile, and L-Leu).

Leucine and isoleucine are isomers, so their mobilograms are presented together. Except for Leu and Ile, all other AAs can be differentiated by their *m/z*. Take the example of the closest isobars of Gln and Lys among all investigated AAs, the *m/z* difference of about 0.036 was readily resolved by the quadrupole time-of-flight (qTOF) analyzer at a resolving power of 44312 at *m/z* 443.126 (Fig.3B). However, D/L-Leu and D/L-Ile are positional isomers that have exactly the same *m/z*. Therefore, unlike other extracted *m/z* windows where only two chiral molecules need to be separated, all four structurally similar chiral molecules need to be separated simultaneously. As presented in Fig.3C, four peaks are recognized respectively as D-Ile, D-Leu, L-Ile, and L-Leu according to the mobility distributions of their pure standards in Fig.S1. The above results demonstrated the powerful ability of FDAA derivatization-enabled differentiation of AAs isomers and enantiomers by IMS, which is important for biological samples since isomers are prevalent in metabolites and should be excluded from target analytes to prevent analysis results from interference. In addition, the method is not dependent on multiple reaction monitoring (MRM) acquisition and therefore has broader instrument applicability. Furthermore, considering the use of most AAs down to 87.7 nM in the mixture, the high enantiomeric resolution, selectivity, and sensitivity of the FDAA derivatization has been demonstrated, which drove us to apply this method to the analysis of real samples.

### Analysis of chiral amino acids in brain extracts

Although the importance of D-amino acids in mammals has been widely recognized, such as the coagonist role of D-Ser at the NMDA receptors^3,48^, the function of most D-AAs is still not completely clear and requires more powerful separation methods for further investigation. In this study, the mouse brain was used as a biological model to explore the utility of FDAA derivatization for chiral IMS analysis. The EIMs of all observed amino acid derivatizes in brain extracts are presented in Fig.S5. In general, 15 amino acids were identified by comparing their *m/z* and mobility values with pure standards after FDAA derivatization. As a representative of Ala shown in Fig.S6, FDAA derivatized Ala exhibits a peak at *m/z* 386.069 in the mass spectrum of mouse brain extracts. In the EIMs of *m/z* 386.069±0.01, an interference peak (grey) is observed and excluded from the FDAA-Ala species by comparing the mobility values with the FDAA-Ala standard. For Tyr and Trp, they were not counted in the 15 identified amino acids due to the *m/z* overlap from interference peaks, but the derivatized analytes can be recognized in their EIMs, as shown in Fig.S7. However, Asp and Glu are not identified owing to their acidic side chains leading to poor ionization efficiency in the positive mode electrospray ionization (ESI).

The quantitative determination of enantiomeric ratio (*er*) in biological samples is of great importance since *er* will change at different development stages of living organisms. The *er* of Ser has been demonstrated to decrease in the mouse brain with increasing age^49^. Ser and Met were chosen as examples to determine *er* in the mouse brain extracts (Fig.4). FDAA-AA standard mixtures with varying concentration ratios of D/L were used to obtain the linear calibration curves between the area ratio of D/L (*r*) and *er* in Fig. 4A and 4B. Good linear curves were realized for Ser and Met with D-AA down to 0.5% in the AA mixture, demonstrating the utility of FDAA derivatization for *er* determination in real samples. The *er* of Ser and Met in mouse brain extracts was determined and is shown in Fig.4C and 4D. The *er* of Ser of about 30% is similar to the D-Ser % measured by LC in mice cortex^50^, but the *er* of Met has not been reported yet. In summary, FDAA derivatization is demonstrated as a powerful method for chiral amino acid separation by TIMS-MS platform.

**Fig.4.**
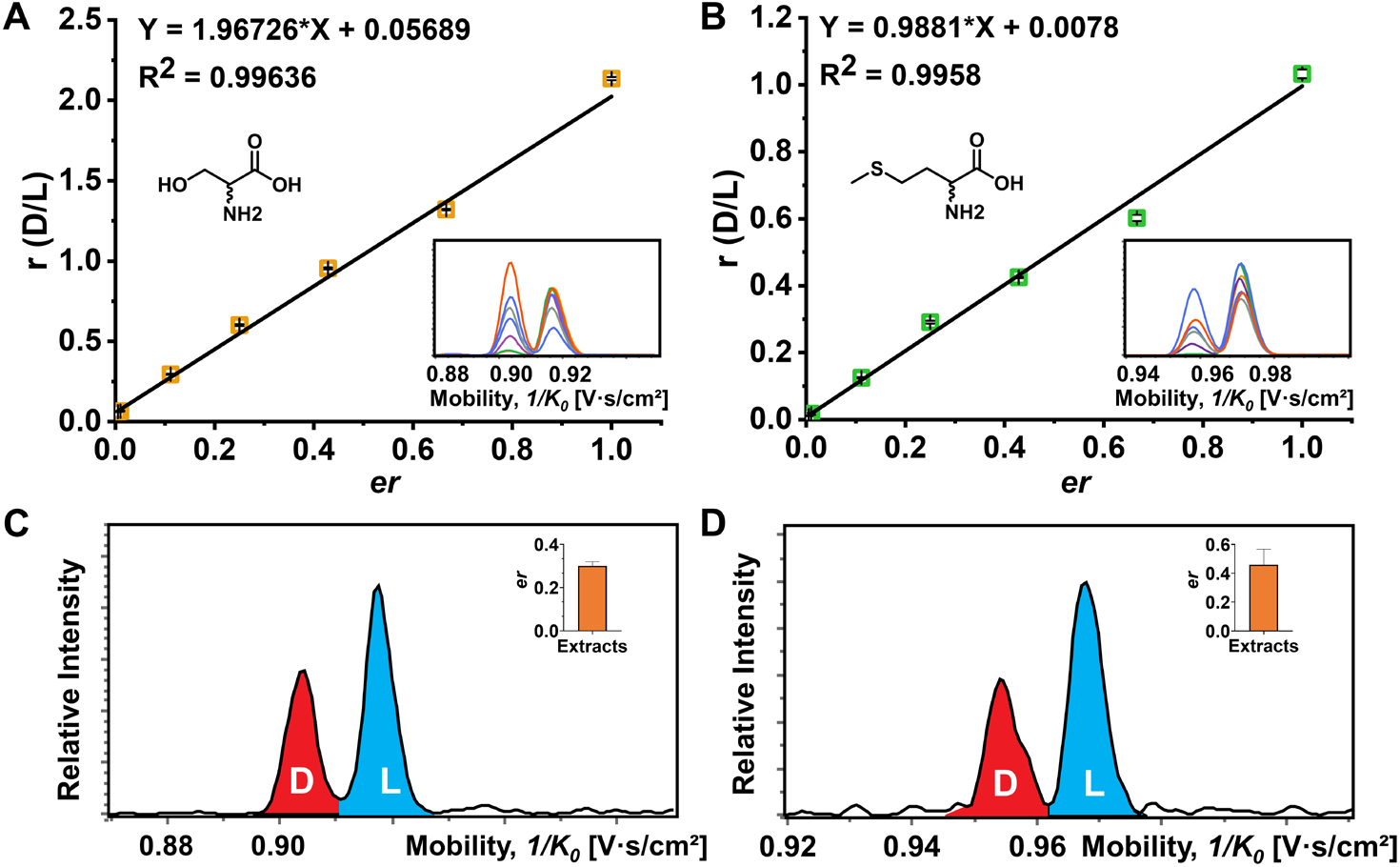
Determination of enantiomeric ratio (*er*) of Ser and Met. (A) The plot of peak area ratio against enantiomeric ratio (D/L) of Ser. (B) The plot of peak area ratio against enantiomeric ratio (D/L) of Met. The corresponding extracted ion mobilograms of D/L-Ser or DL-Met at different mixing ratios are embedded. Four technical replications of the TIMS-MS analysis were performed. (C) Extracted ion mobilograms and the corresponding *er* of Ser (inserted bar chart) of the mouse brain extracts. (D) Extracted ion mobilograms and the corresponding *er* of Met (inserted bar chart) of the mouse brain extracts. The TIMS-MS analysis for mouse brain extracts was carried out six times.

### Visualization of chiral amino acids

Recently, mass spectrometry imaging has been used to reveal the spatial distribution of isomeric metabolites in biological systems^51–56^. Based on the FDAA-enabled chiral separation ability exhibited in the tissue extracts, we further investigate the separation ability of FDAA derivatization in tissue section by MALDI (matrix-assisted laser desorption/ionization) TIMS MSI, 38 AAs were directly deposited on the right half of the mouse brain section, and MSI was performed on the rectangular area covering the whole mouse brain section. In the mass spectrum shown in Fig.5A, all *m/z* of [FDAA-AA + 2Na - H]^+^ ions are detected, demonstrating the success of the on-tissue FDAA reaction, but *m/z* 418.040 was excluded from Cys by comparing its EIMs with Cys standards in Fig.S8, possibly due to the easily oxidizable properties of Cys. Each pair of the remaining AAs were separated by MALDI TIMS-MS and the two-dimension features (*m/z* vs. mobility) of the 18 pair AAs with defined windows were extracted and drawn in Fig.S9, demonstrating the separation capacity for chiral AAs after FDAA derivatization by TIMS-MS even in long-term MALDI MSI acquisition. The ion images of all separated chiral AAs are shown in Fig.5B. For most AAs, higher ion signals can be observed in outside areas compared to the brain part, possibly due to the presence of competing molecules in the tissue region. L-Gln, D/L-Asp and D/L-Glu show the opposite trend since these amino acids have a higher abundance than other AAs in the brain tissue^50^ and present relatively clear ion images in the undeposited left half of the brain section. According to the results of standard addition method, endogenous chiral amino acids can be identified and visualized in Fig.S10. Overall, these results validate the applicability of the developed method to differentiate and visualize chiral amino acids in biological samples using MALDI TIMS MSI.

**Fig.5.**
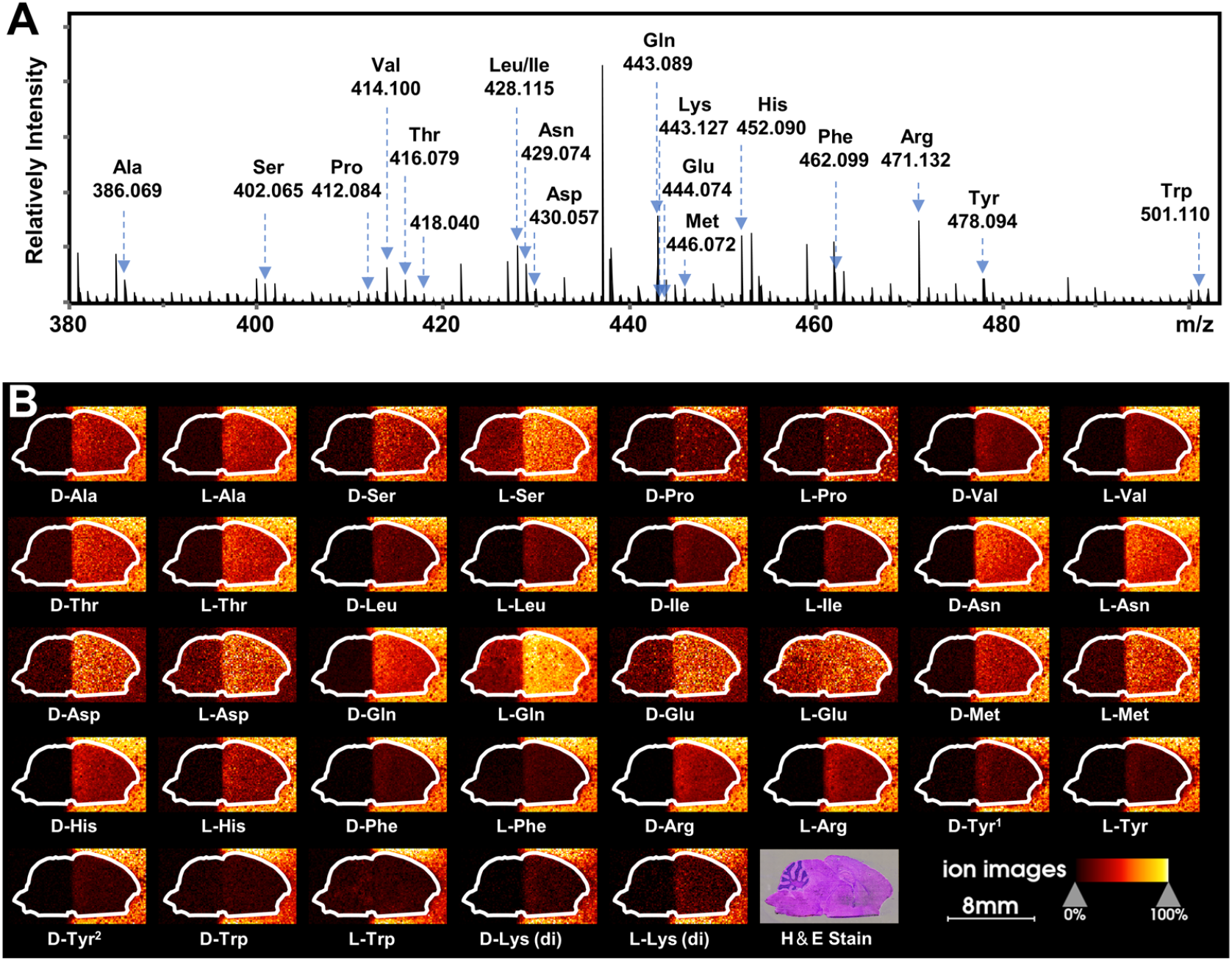
MALDI TIMS MSI of chiral amino acids in mouse brain tissue after the deposition of 38 AA standards by on-tissue FDAA derivatization. (A) MALDI mass spectrum recorded from the section of mouse brain; amino acid derivative is annotated as AA based on accurate mass and mobility alignments. (B) ion images of detected chiral AAs. The extracting windows of *m/z* and mobility were set according to Fig.S9†.

## Conclusions

In this study, A powerful TIMS-based method was developed for chiral amino acid analysis using FDAA derivatization. Chiral separation was achieved for all 19 pairs of proteinogenic amino acids labeled with FDAA using the same ionic form, avoiding the optimization of reference compounds and metal ions for specific analytes. The high selectivity and sensitivity of FDAA combined with TIMS-MS for chiral discrimination were also demonstrated in the simultaneous separation of the mixture of 38 AAs (19 pairs of DL-forms) down to the nM concentration range. Using the FDAA derivatization method, we detected 15 AAs simultaneously and excluded isomeric and isobaric interference in mouse brain extracts. As a proof of concept, *er* was determined for Ser and Met in the brain extracts. The separation capacity of FDAA derivatization was also proved in MALDI TIMS MSI of the mouse brain section with the deposition of 38 AAs. In general, the developed method shows powerful potential for IMS-based separation of chiral amino acids both in ESI analysis and MALDI MSI, which widened the path to the further investigation of biological functions and mechanisms of chiral molecules in complex biological samples. In addition, the effectiveness of this CDA for other metabolites containing amine group is also well worth studying to expand potential IMS-based applications.

## Acknowledgments

This work was supported by The Shenzhen Science and Technology Innovation Commission (2021Szvup134), SKLEBA Research Grant (SKLP_2021_P04), and the Start-up Grant from Hong Kong Baptist University.

