## Supplemental tables and figures for "Chiral Derivatization-enabled Discrimination and Visualization of Proteinogenic Amino Acids by Ion Mobility Mass Spectrometry"

Electronic Supporting Information

**Fig. S10.** MALDI TIMS MSI of endogenous chiral amino acids (glutamic acid) in a mouse brain tissue section .......... 16

### 1.1 Animal preparation

C57Bl/6 mice were obtained from Laboratory Animal Service Centre at The Chinese University of Hong Kong (Hong Kong SAR, China). The mice were housed in standard conditions with a 12h light /12h dark cycle and fed with sterilized water and standard laboratory feed. The mice were sacrificed, and brain tissues were dissected and stored at -80 ℃ prior to metabolites extraction and section preparation. Experimental protocols were approved by the Hong Kong Baptist University Committee on the Use of Human and Animal Subjects in Teaching and Research.

### 1.2 Amino acids extraction from mouse brain tissue

The preparation of mouse brain extracts followed the workflow previously reported[1].50 mg of brain tissue samples were homogenized in 500 $\mu$L of ice-cold MeOH/$H_2O$ (4:1, v: v) using Bullet Blender Storm 24 (Next Advance, Troy, NY, USA). 300 $\mu$L of homogenate was extracted and mixed with 180 $\mu$L of chloroform and 60 $\mu$L of $H_2O$. The mixture was vortexed for 1 min, rested for 5 mins to equilibrate to room temperature, and then centrifuged at 20 000 g for 10 mins at 4 ℃. The up layer was collected, transferred to a 1.5 mL microcentrifuge tube, and stored at -80 ℃ prior to the FDAA derivatization reaction.

### 1.3 Sample preparation for MALDI MSI

Mouse brain tissue was cryosectioned at 10 μm using CryoStar Nx70 cryostat (Thermal Fisher Scientific, Walldorf, Germany) at a chamber temperature of -20 °C. The tissue cryosections were then transferred onto the conductive side of indium tin oxide (ITO)-coated glass slides (2.5 cm x 7.5 cm, Delta Technologies, Loveland, CO, USA). The tissue sections were rinsed with cold chloroform for 15s at -18 °C to remove lipids from the tissue section, which was reported to improve the detection of small molecule metabolites[2]. Adjacent tissue slides were stained with hematoxylin and eosin (H&E staining), and optical images were captured using Confocal FLIM Imaging System (Nikon C2si Plus, Japan).

The mixing solution of 38 AAs (About 212 $\mu$M of each AA) was prepared in an aqueous $NaHCO_3$ (100 mM) solution and then was diluted twice using MeOH. Half of the brain section was covered with a coverslip (Corning, NY, USA) to prevent standards from depositing on this half. The AA mixture was applied to the tissue section covering the dimensions of 2.6 cm x 3 cm using a custom-built matrix deposition system. The setup of the spray system is similar to our previously reported electrospray deposition device[3]. The AA mixture was delivered by a springe pump (Chemyx Fusion 101, Chemyx, Stafford, TX, USA) with a flow rate of 10 $\mu$L/min for 30mins through a 100 μm i.d. quartz capillary (IDEX Corp., Northbrook, IL, USA) to the spray nozzle with $N_2$ sheath gas (99.995 %) of 70 psi heated to 50 °C. The spraying voltage was set to +5 kV using a high voltage power supply (PS 350, Stanford Research Systems, Sunnyvale, CA, USA). The motorized stage was set at a velocity of 1098 mm/min with 1 mm track spacing.

On-tissue FDAA derivatization (Fig. 5A) was performed on the AA-deposited slide for 60 mins using the deposition system. FDAA (20 mM) in a MeOH/THF (1:1, v/v) solution and an aqueous $NaHCO_3$ (100 mM) solution were mixed and delivered at a flow rate of 30 $\mu$L/min The sheath gas was decreased to 35 psi and the spraying dimensions were set to 2.6 cm x 3 cm to ensure a 0.5 cm spacing at least between the spraying edge and the tissue on each side.

After derivatization, a DHAP solution (5 mg/mL) in MeOH was applied onto the tissue section using the same settings as the deposition of AAs, except that the thickness and dimensions were changed to 13 cycles and 2.6 cm x 3 cm, respectively.

**Table S1.** Parameters of direct-infusion TIMS-MS in Tune

| | | | |
|---|---|---|---|
| Transfer | | | |
| Funnel 1 RF | 350.0 $V_{pp}$ | isCID Energy | 0.0 eV |
| Funnel 2 RF | 350.0 $V_{pp}$ | Deflection Delta | 70.0 V |
| Multipole RF | 350.0 $V_{pp}$ | | |
| Quadrupole | | | |
| Ion Energy | 5.0 eV | Low Mass | 300.00 *m/z* |
| Collision Cell | | | |
| Collision Energy | 10.0 eV | Collision RF | 1800.0 $V_{pp}$ |
| Focus Pre TOF | | | |
| Transfer Time | 60.0 $\mu$s | Pre Pulse Storage | 8.0 $\mu$s |

**Table S2.** Parameters of MALDI TIMS-MS in Tune

| | | | |
|---|---|---|---|
| Transfer | | | |
| MALDI Plate Offset | 50.0 V | Deflection 1 Delta | 70.0 V |
| Funnel 1 RF | 200.0 $V_{pp}$ | isCID Energy | 0.0 eV |
| Funnel 2 RF | 220.0 $V_{pp}$ | Multipole RF | 220.0 $V_{pp}$ |
| Quadrupole | | | |
| Ion Energy | 5.0 eV | Low Mass | 150.00 m/z |
| Collision Cell | | | |
| Collision Energy | 10.0 eV | Collision RF | 1500.0 $V_{pp}$ |
| Focus Pre TOF | | | |
| Transfer Time | 60.0 $\mu$s | Pre Pulse Storage | 5.0 $\mu$s |

**Table S3.** Elution orders and $R$ of FDAA-AA diastereomers obtained from TIMS-MS

| FDAA-AA | First eluting AA | $R_D$ | $R_L$ |
|---|---|---|---|
| Ala | D | 123.3 | 147.5 |
| Arg | D | 146.7 | 134.4 |
| Asn | D | 112.7 | 148.1 |
| Asp | D | 93.8 | 168.4 |
| Cys[a] | L | 184.9 | 114.4 |
| Gln | D | 165.4 | 164.9 |
| Glu | D | 122.4 | 146.9 |
| His | D | 176.5 | 172.0 |
| Leu | D | 144.3 | 164.6 |
| Ile | D | 146.5 | 164.6 |
| Lys | D | 150.3 | 153.8 |
| Met | D | 140.9 | 147.4 |
| Phe | D | 157.9 | 134.0 |
| Pro | D | 140.0 | 181.4 |
| Ser | D | 137.4 | 141.5 |
| Thr | D | 157.8 | 160.8 |
| Trp | D | 154.0 | 163.5 |
| Tyr | D | 163.5 | 107.1 |
| Val | D | 172.7 | 132.4 |

[a]bis-derivatives

**Table S4.** Extracting windows of m/z and mobility in Fig. S10

| FDAA-AA | m/z up limit | m/z lower limit | $1/k_0$ up limit | $1/k_0$ lower limit |
|---|---|---|---|---|
| D-Ala | 386.059 | 386.079 | 0.8960 | 0.9080 |
| L-Ala | 386.059 | 386.079 | 0.9120 | 0.9260 |
| D-Arg | 471.122 | 471.142 | 0.9895 | 0.9975 |
| L-Arg | 471.122 | 471.142 | 0.9985 | 1.0065 |
| D-Asn | 429.064 | 429.084 | 0.9265 | 0.9365 |
| L-Asn | 429.064 | 429.084 | 0.9380 | 0.9480 |
| D-Asp | 430.048 | 430.068 | 0.9200 | 0.9360 |
| L-Asp | 430.048 | 430.068 | 0.9400 | 0.9520 |
| D-Gln | 443.079 | 443.099 | 0.9460 | 0.9560 |
| L-Gln | 443.079 | 443.099 | 0.9580 | 0.9680 |
| D-Glu | 444.063 | 444.083 | 0.9500 | 0.9580 |
| L-Glu | 444.063 | 444.083 | 0.9610 | 0.9670 |
| D-His | 452.080 | 452.100 | 0.9580 | 0.9660 |
| L-His | 452.080 | 452.100 | 0.9810 | 0.9880 |
| D-Leu | 428.105 | 428.125 | 0.9700 | 0.9706 |
| L-Leu | 428.105 | 428.125 | 0.9830 | 0.9890 |
| D-Ile | 428.105 | 428.125 | 0.9580 | 0.9680 |
| L-Ile | 428.105 | 428.125 | 0.9770 | 0.9810 |
| D-Lys (di) | 695.164 | 695.184 | 1.1275 | 1.1415 |
| L-Lys (di) | 695.164 | 695.184 | 1.1500 | 1.1640 |
| D-Met | 446.062 | 446.082 | 0.9450 | 0.9590 |
| L-Met | 446.062 | 446.082 | 0.9650 | 0.9790 |
| D-Phe | 462.089 | 462.109 | 0.9620 | 0.9740 |
| L-Phe | 462.089 | 462.109 | 0.9760 | 0.9900 |
| D-Pro | 412.074 | 412.094 | 0.9140 | 0.9240 |
| L-Pro | 412.074 | 412.094 | 0.9320 | 0.9440 |
| D-Ser | 402.055 | 402.075 | 0.8980 | 0.9100 |
| L-Ser | 402.055 | 402.075 | 0.9120 | 0.9240 |
| D-Thr | 416.069 | 416.089 | 0.9220 | 0.9320 |
| L-Thr | 416.069 | 416.089 | 0.9320 | 0.9440 |
| D-Trp | 501.100 | 501.120 | 1.0060 | 1.0200 |
| L-Trp | 501.100 | 501.120 | 1.0200 | 1.0340 |
| D-Try[1] | 478.084 | 478.104 | 0.9860 | 0.9980 |
| L-Try | 478.084 | 478.104 | 1.0000 | 1.0140 |
| D-Try[2] | 478.084 | 478.104 | 1.0280 | 1.0420 |
| D-Val | 414.090 | 414.110 | 0.9350 | 0.9450 |
| L-Val | 414.090 | 414.110 | 0.9500 | 0.9620 |

di: bis-derivatives

[1]first eluted mobility peak of D-Try

[2]second eluted mobility peak of D-Try

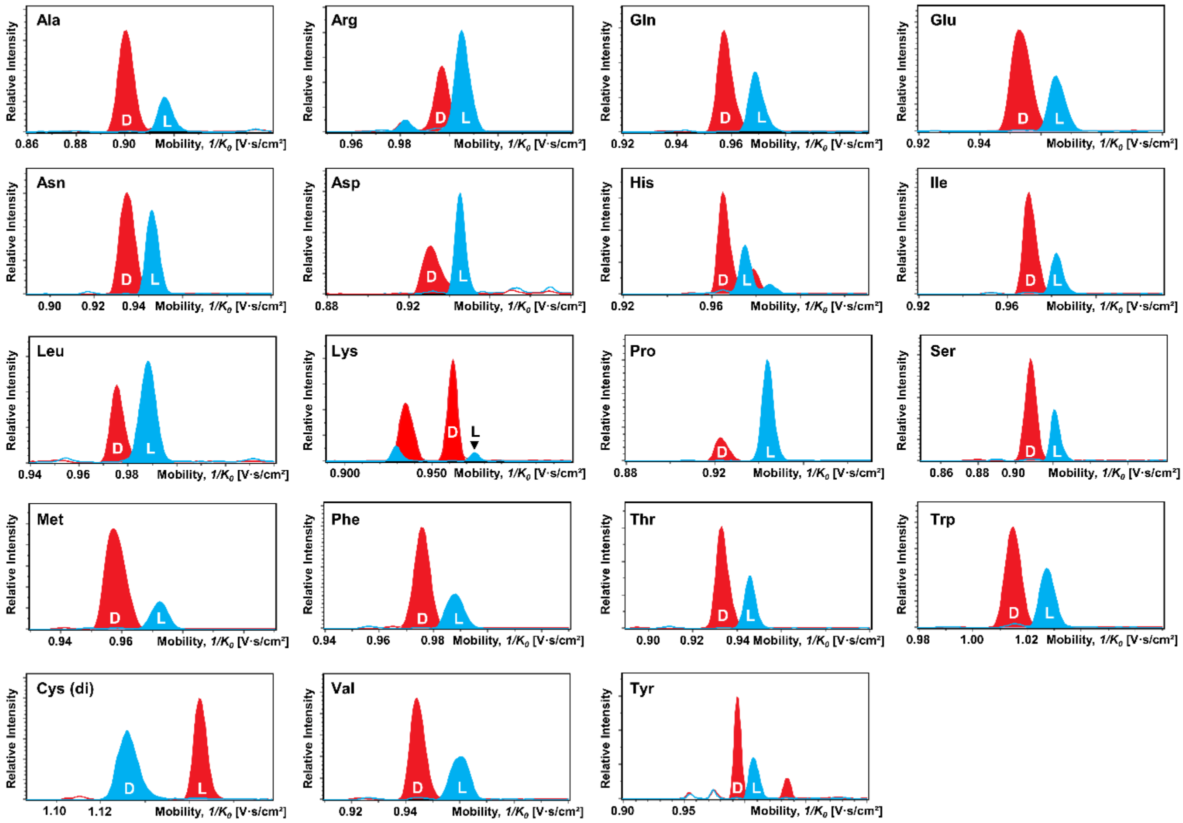

**Fig. S1.** Overlay EIMs of a single chiral amino acid standard after FDAA derivatization. The corresponding D-forms and L-forms were measured by TIMS-MS separately.

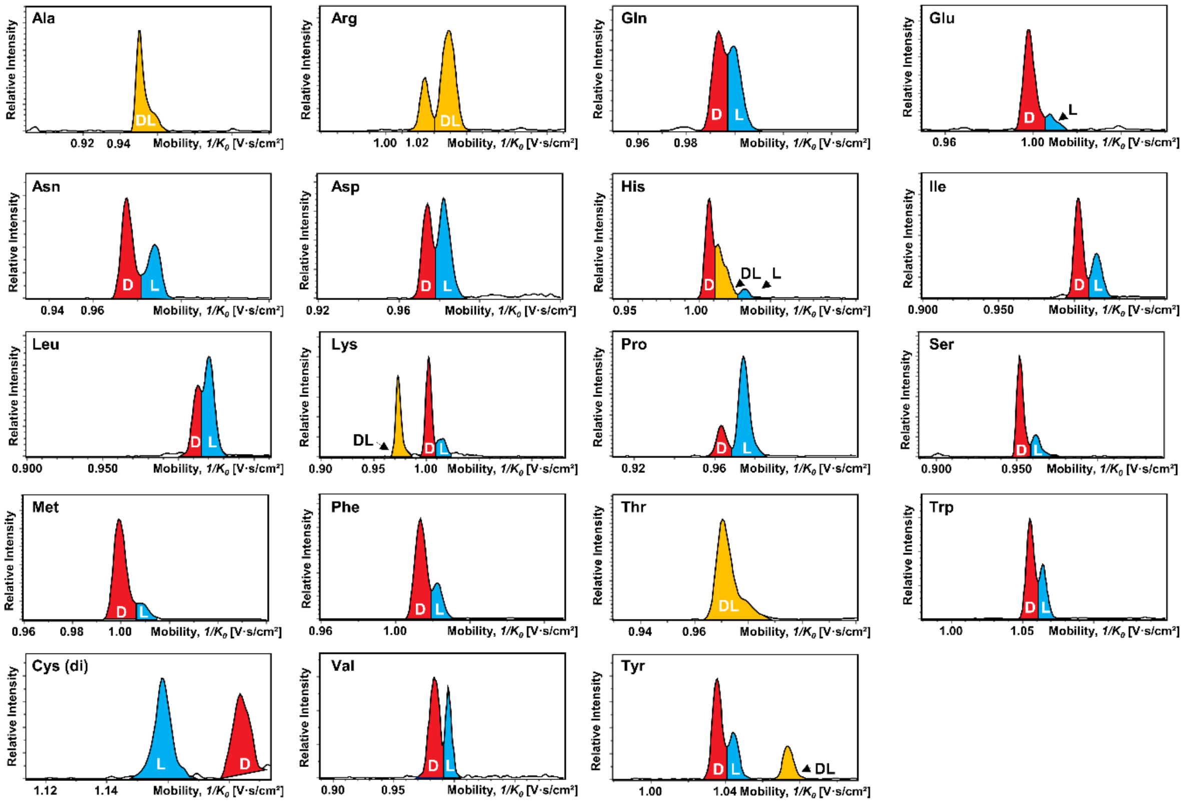

**Fig. S2.** EIMs of 19 pairs of proteinogenic AA (D:L = 1:1) standards after FDVA derivatization.

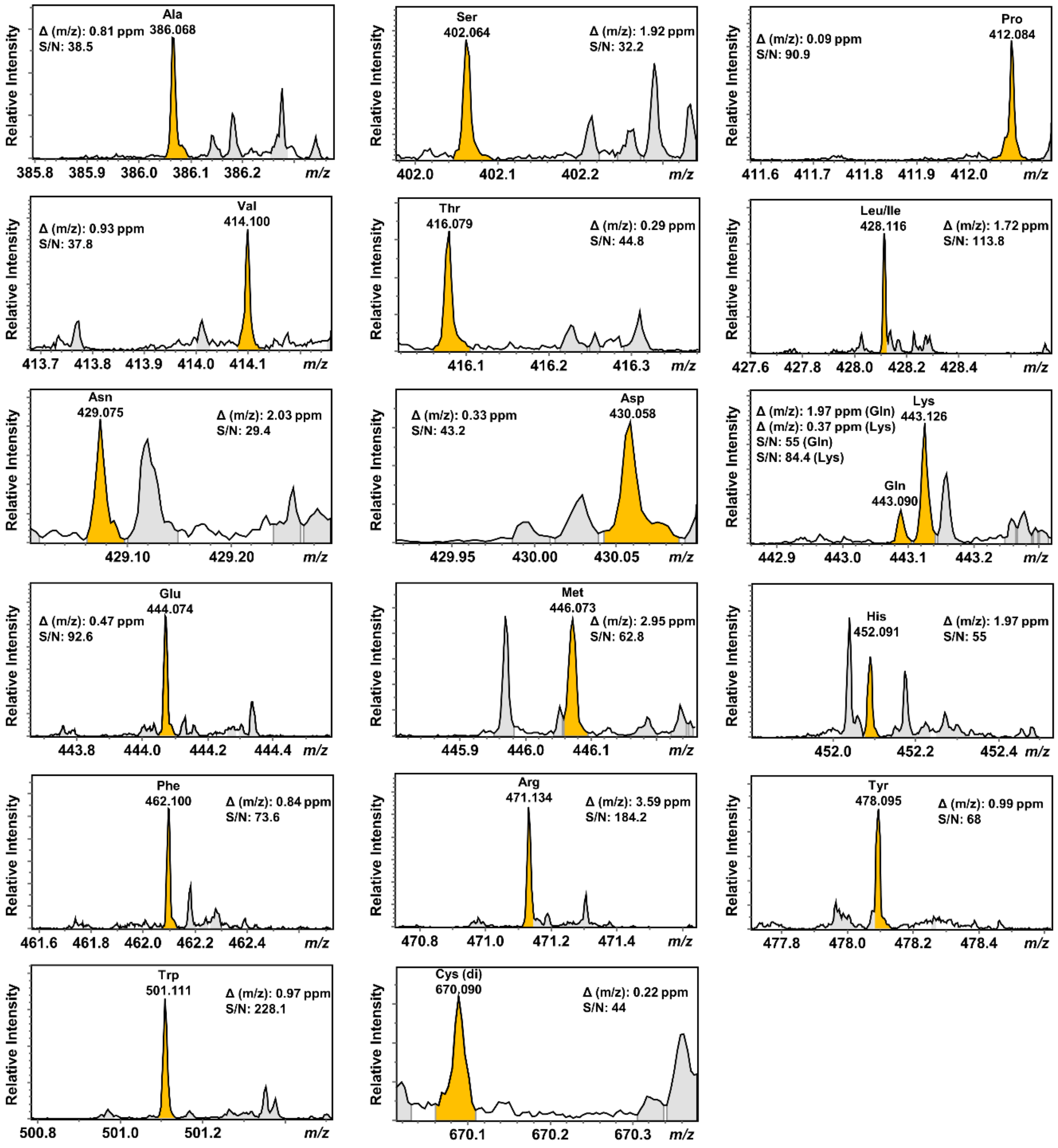

**Fig. S3.** Mass spectra of 38 mixed amino acids corresponding to Fig. 3. The mass deviation and signal to noise ratio (S/N) of highlighted peaks are presented as well.

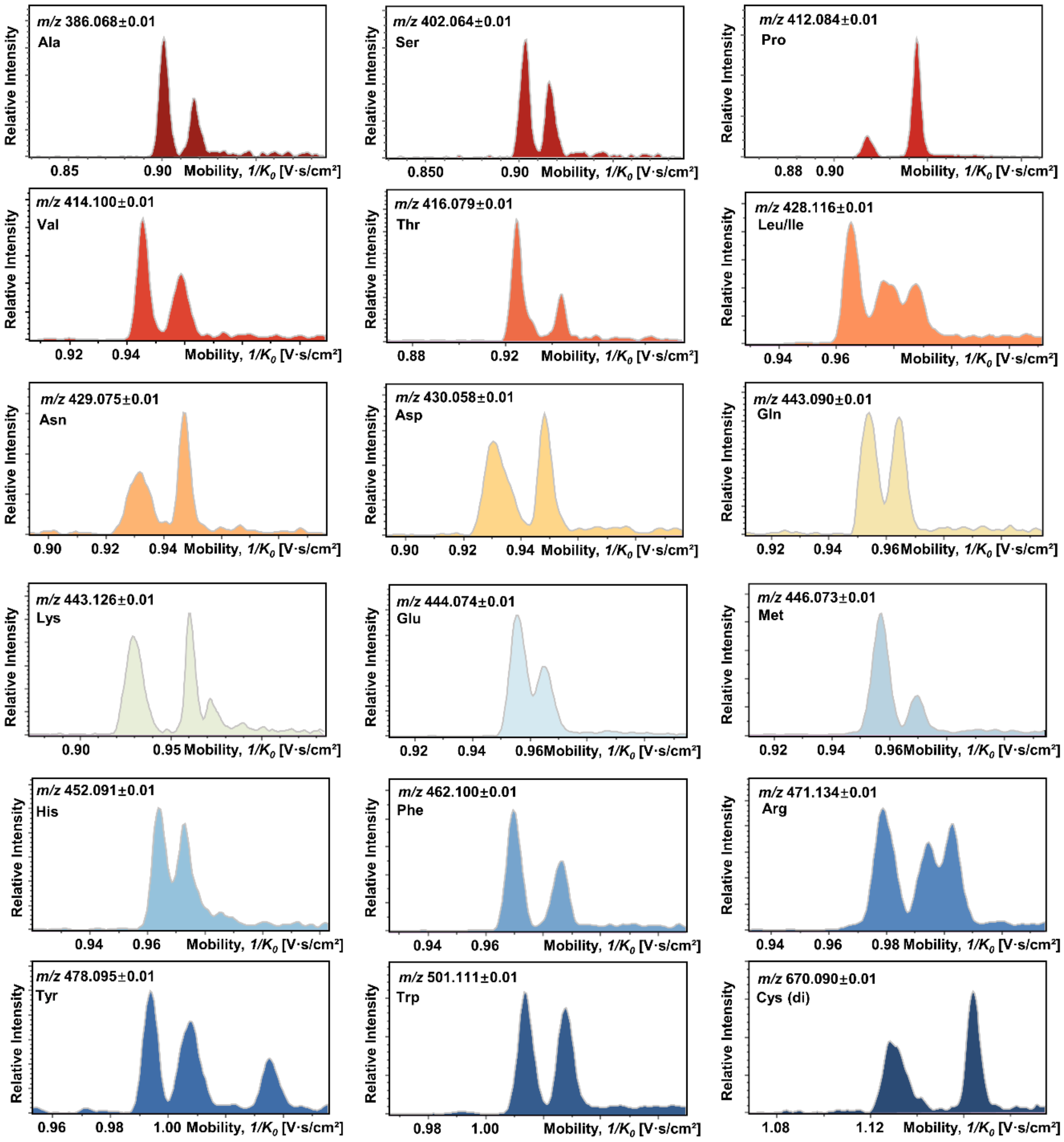

**Fig. S4.** Separately listed EIMs of the mixture of 38 chiral AAs corresponding to Fig. 3.

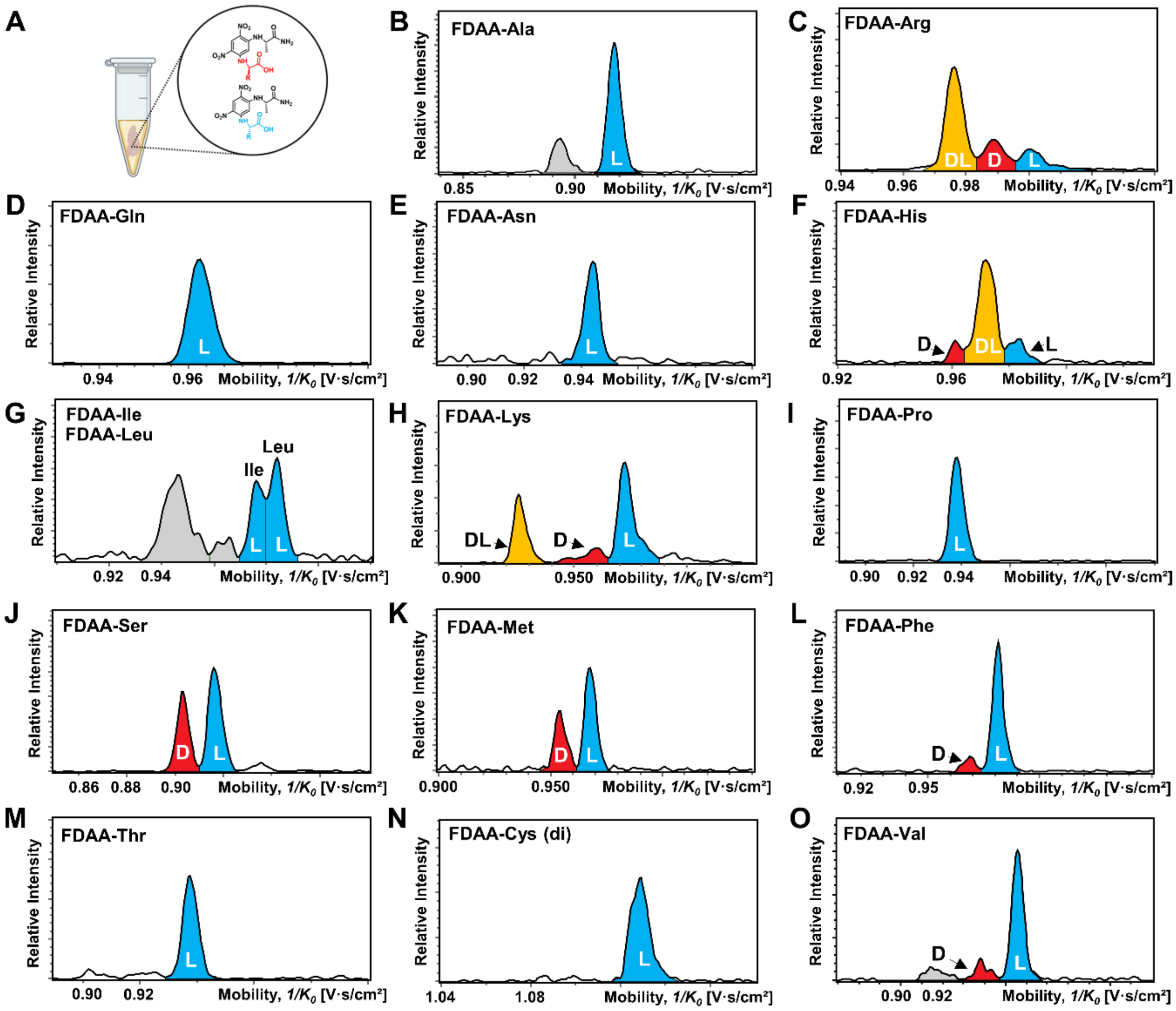

**Fig. S5.** EIMs of 15 AAs measured in the mouse brain extracts after FDAA derivatization. (A) Schematic diagram of mouse brain extracts, which was created with BioRender.com. (B-O) EIMs of FDAA-AA.

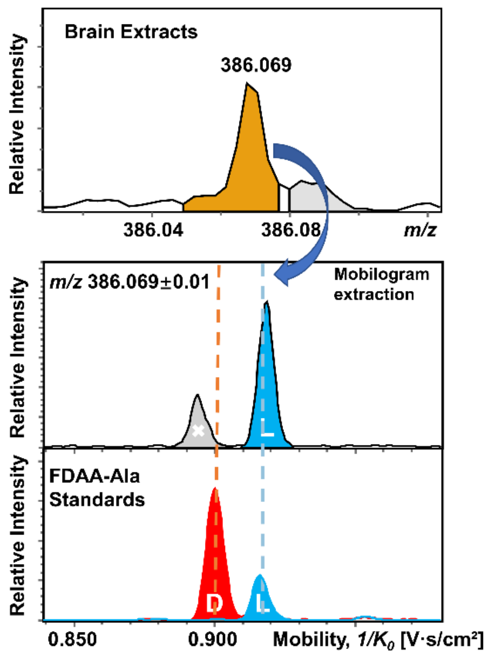

**Fig. S6.** An example of amino acid (Ala) identification in mouse brain extracts using FDAA derivatization by TIMS-MS. Mobility distribution in the EIM of m/z 386.069 ± 0.01 is compared with FDAA-Ala pure standard. Peaks in red, blue, and grey traces represent D-AA, L-AA, and interference ions, respectively.

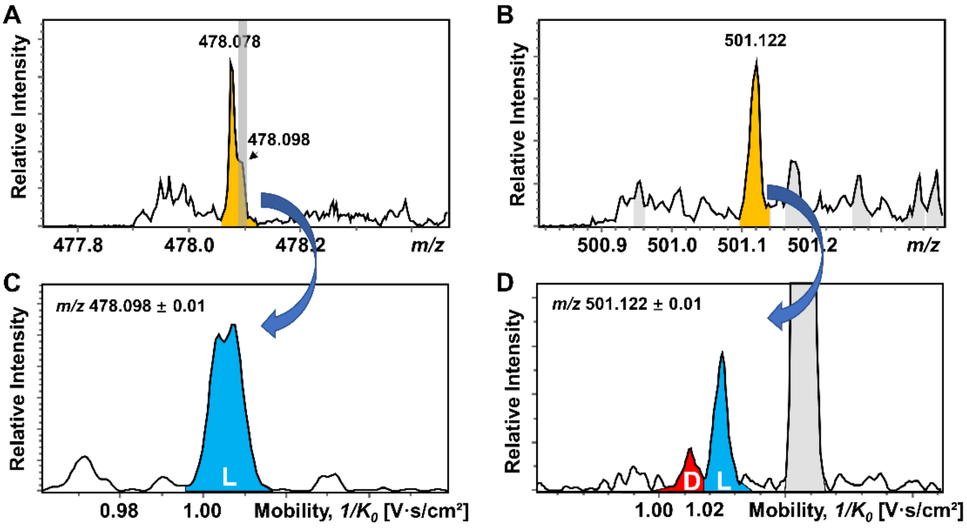

**Fig. S7.** Mass spectra and EIMs of Tyr and Trp measured in the mouse brain extracts after FDAA derivatization. (A) Mass spectrum corresponding to FDAA-Tyr. (B) Mass spectrum corresponding to FDAA-Trp. (C) EIMs of *m/z* 478.098 ± 0.01. (D) EIMs of *m/z* 501.122 ± 0.01. Peaks in red, blue, and grey traces represent D-AA, L-AA, and interference ions, respectively. While the m/z of target analytes are overlapped with an interference peak, the target analytes can be recognized in the EIMs.

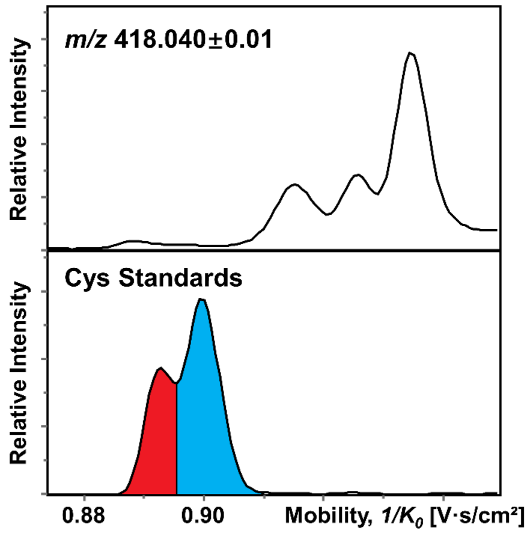

**Fig. S8.** EIMs of m/z 418.040 ± 0.01 measured in the mouse brain section and Cys standards after FDAA derivatization.

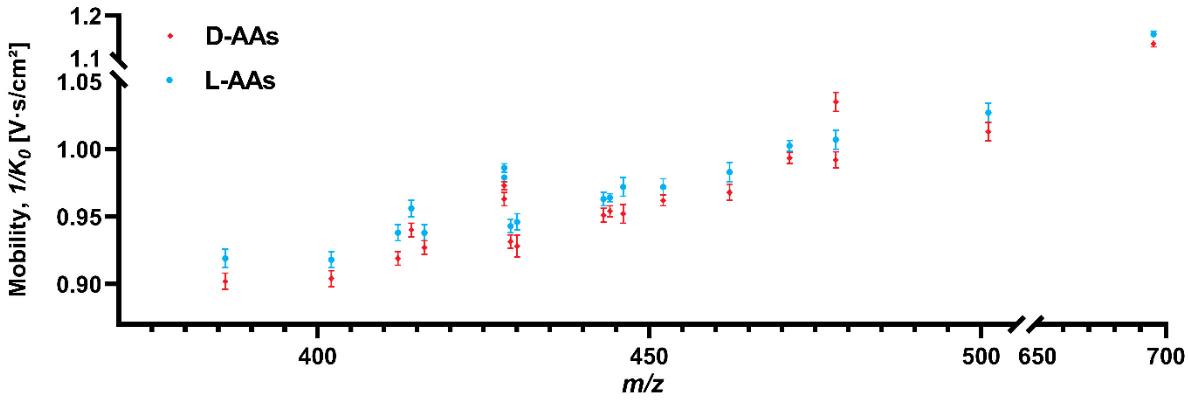

**Fig. S9.** Extracting windows of m/z and mobility in Fig. 5. Detailed values are listed in Table S4. The windows are set based on that no mutual interference occurs for ion images.

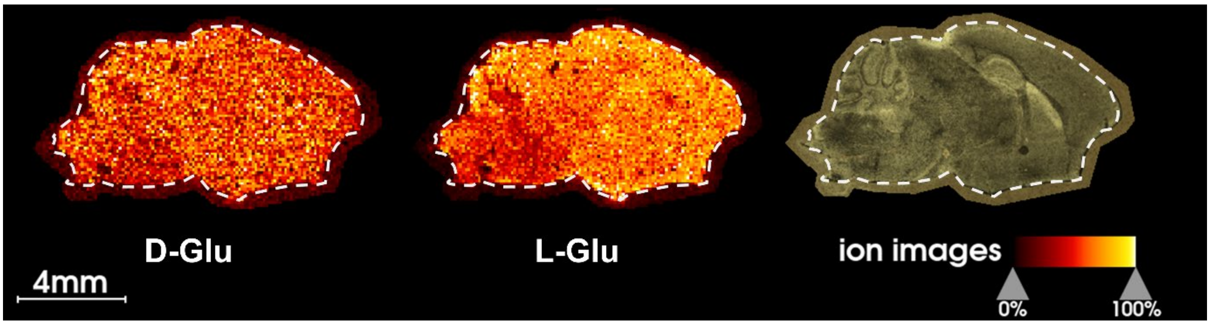

**Fig. S10.** MALDI TIMS MSI of endogenous chiral amino acids (glutamic acid) in a mouse brain tissue section.
